# Whole-brain mapping in adult zebrafish and identification of a novel tank test functional connectome

**DOI:** 10.1101/2024.08.16.607981

**Authors:** Neha Rajput, Kush Parikh, Ada Squires, Kailyn K. Fields, Matheu Wong, Dea Kanani, Justin W. Kenney

## Abstract

Identifying general principles of brain function requires the study of structure-function relationships in a variety of species. Zebrafish have recently gained prominence as a model organism in neuroscience, yielding important insights into vertebrate brain function. Although methods have been developed for mapping neural activity in larval animals, we lack similar techniques for adult zebrafish that have the advantage of a fully developed neuroanatomy and larger behavioral repertoire. Here, we describe a pipeline built around open-source tools for whole-brain activity mapping in freely swimming adult zebrafish. Our pipeline combines recent advances in histology, microscopy, and machine learning to capture *cfos* activity across the entirety of the adult brain. Images captured using light-sheet microscopy are registered to the recently created adult zebrafish brain atlas (AZBA) for automated segmentation using advanced normalization tools (ANTs). We used our pipeline to measure brain activity after zebrafish were subject to the novel tank test. We found that *cfos* levels peaked 15 minutes following behavior and that several regions containing serotoninergic, dopaminergic, noradrenergic, and cholinergic neurons were active during exploration. Finally, we generated a novel tank test functional connectome. Functional network analysis revealed that several regions of the medial ventral telencephalon form a cohesive sub-network during exploration. We also found that the anterior portion of the parvocellular preoptic nucleus (PPa) serves as a key connection between the ventral telencephalon and many other parts of the brain. Taken together, our work enables whole-brain activity mapping in adult zebrafish for the first time while providing insight into neural basis for the novel tank test.

## Introduction

A fundamental goal of neuroscience is to understand how patterns of brain activity give rise to behavior. Identifying general principles of brain function is facilitated by cross species comparisons. Over the past two decades, zebrafish have started contributing to our understanding of the brain, a trend that promises to continue due to their low cost, ease of genetic manipulation, and sophisticated behavioral repertoire (Gerlai, 2023; Kenney, 2020; Loring et al., 2020). Although several methods have been developed for whole-brain activity mapping in larval zebrafish (Ahrens et al., 2012; Portugues et al., 2014; Randlett et al., 2015; Shainer et al., 2023), equivalent approaches have yet to be developed for adult stage animals.

Adult and larval zebrafish each have distinct advantages and disadvantages in the study of brain-behavior relationships. Whereas larval animals are amenable to high throughput work due to their small size and transparency, adults have the advantage of mature neuroanatomy and more extensive behavioral repertoire. This behavioral repertoire includes a wide variety of social behaviors (Gerlai, 2014; Jones and Norton, 2015; Kareklas et al., 2023), short and long-term associative, non-associative, and spatial memories (Gerlai, 2020; Kenney, 2020), and different types of exploratory behaviors (Cachat et al., 2010; Rajput et al., 2022; Toms and Echevarria, 2014). Thus, to fully realize the utility of zebrafish as a model organism in neuroscience, methods for whole-brain mapping are also required for adult zebrafish.

Whole-brain activity mapping can yield unexpected insights into brain function that may be lost using more targeted methods. Measuring neural activity across the entire brain also facilitates the use of powerful analytic tools, like network analysis, that captures complex interactions and improves predictions of brain-behavior relationships (Vetere et al., 2017; Wheeler et al., 2013). However, mapping whole-brain activity presents several technical challenges. One roadblock is that the brain of adult animals is not transparent, and thus requires the use of tissue clearing (Richardson et al., 2021). Imaging intact organs presents another technical hurdle due to the increased volume, a challenge met by the recent development of light-sheet microscopy (Hillman et al., 2019). Finally, whole-brain mapping results in large amounts of data that cannot be analyzed via traditional approaches like manual counting and segmentation. We tackled this challenge by combining advances in machine learning to automate cell detection (Tyson et al., 2021) and image registration (Gholipour et al., 2007) with the recently created digital adult zebrafish brain atlas (AZBA) (Kenney et al., 2021). Here, we describe how we have assembled these tools into a pipeline that enables whole-brain activity mapping in adult zebrafish for the first time.

## Results

### Overview of strategy

We begin by giving an overview of our strategy for whole-brain activity mapping (figure 1) before describing the results of each step in more detail. Following behavior, animals are euthanized and heads fixed in 4% paraformaldehyde overnight. Following careful dissection, brains are rendered optically transparent using iDISCO+ (Renier et al., 2016), which we modified to make it compatible with *in situ* hybridization chain reaction (HCR) for the detection of *cfos* mRNA (Choi et al., 2018; Kramer et al., 2018; Kumar et al., 2021). Imaging intact cleared brain tissue was done using light-sheet microscopy. To automatically identify *cfos* positive cells in the brain, we used the open source CellFinder package (Tyson et al., 2021) that is part of the BrainGlobe suite of Python-based software tools (Claudi et al., 2020). Finally, to automatically parcellate the brain into individual regions, we used advanced normalization tools (ANTs; Avants et al., 2009)) to register images to AZBA (Kenney et al., 2021). The final output of our pipeline is a list of *cfos* positive cell counts for each brain region and each animal. This enables the use of a variety of downstream analytic tools, one example that we demonstrate here is functional network analysis.

**Figure 1.**
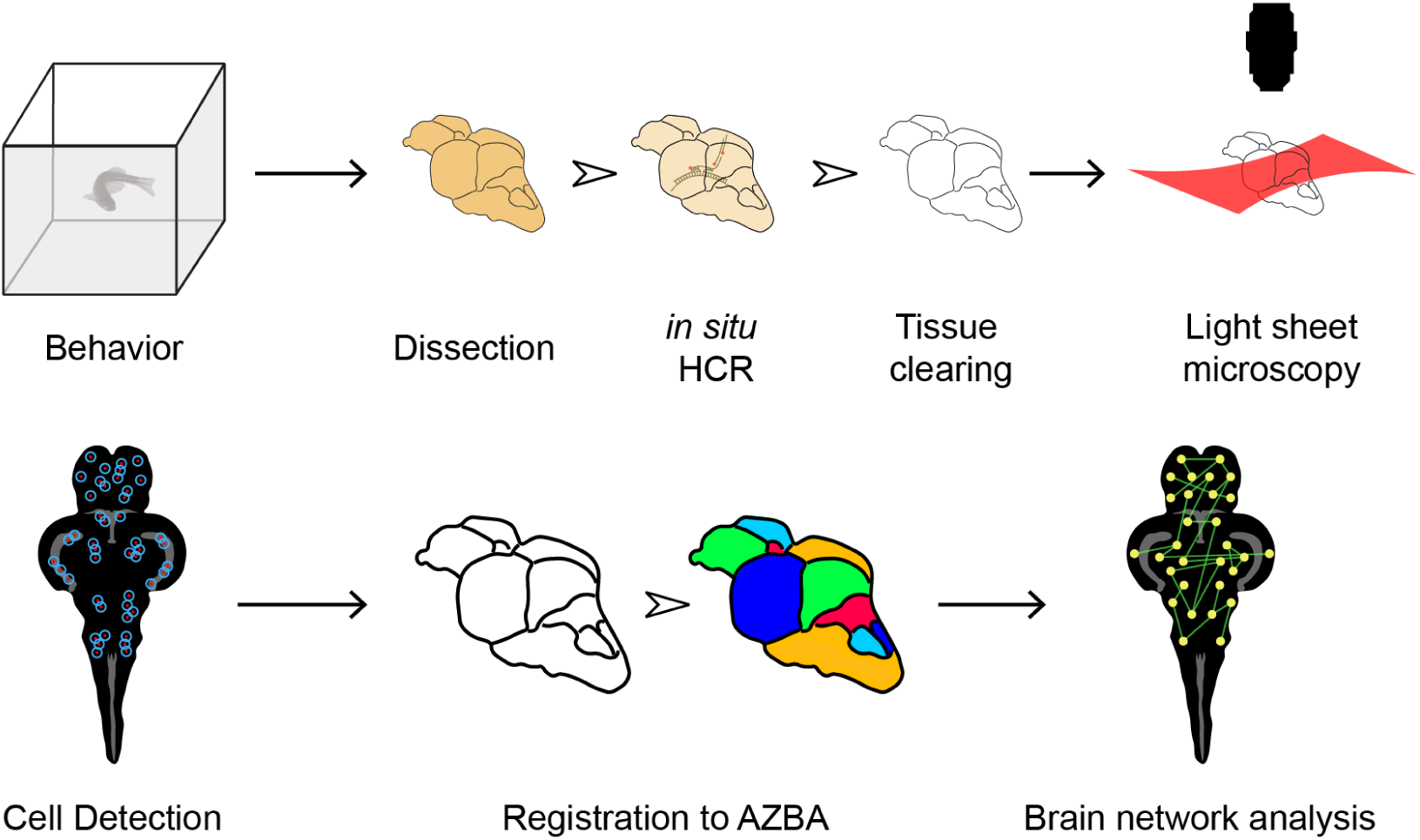
Overview of method for mapping neural activity in adult zebrafish. Following behavior, zebrafish are euthanized and brains carefully removed. *In situ* HCR is then used to label *cfos*. Brians are then cleared using iDISCO and imaged using light-sheet microscopy. Cells are then detected using CellFinder and brains are registered to AZBA. Regional *cfos* counts are then used to generate brain networks for further analysis.

### Automated cell detection

After *in situ* HCR, tissue was cleared using iDISCO+, which allowed us to use light-sheet microscopy to capture whole-brain images in both the *cfos* (Figure 2A, top) and autofluorescence channels (Figure 2A, bottom). Detection of *cfos* positive cells was done using CellFinder (Tyson et al., 2021), an artificial neural net-based supervised machine learning algorithm. The first step in the cell detection process uses image filtering to detect cell shaped objects in the *cfos* image. We found parameters that captured *cfos* positive cells throughout the entire brain (described in the methods section), including areas with cells of different sizes and densities like the telencephalon (Figure 2B) and cerebellum (Figure 2C). Because the cell detection algorithm generated a lot of overlapping cells, we used a custom written Python script to remove cell candidates that were within 9 μm of one another. We then trained the CellFinder artificial neural network by manually labelling 10,597 cells and 7,303 non-cells across five brains. Non-cells were unambiguously identified by the presence of a signal in both the *cfos* and autofluorescence channels, suggesting the presence of background bleeding into the *cfos* channel. Cells only appeared in the *cfos* channel. The resulting network achieved over 95% accuracy where the cells and non-cells were clearly differentiated across several different brain regions (Figure 2B & C).

**Figure 2.**
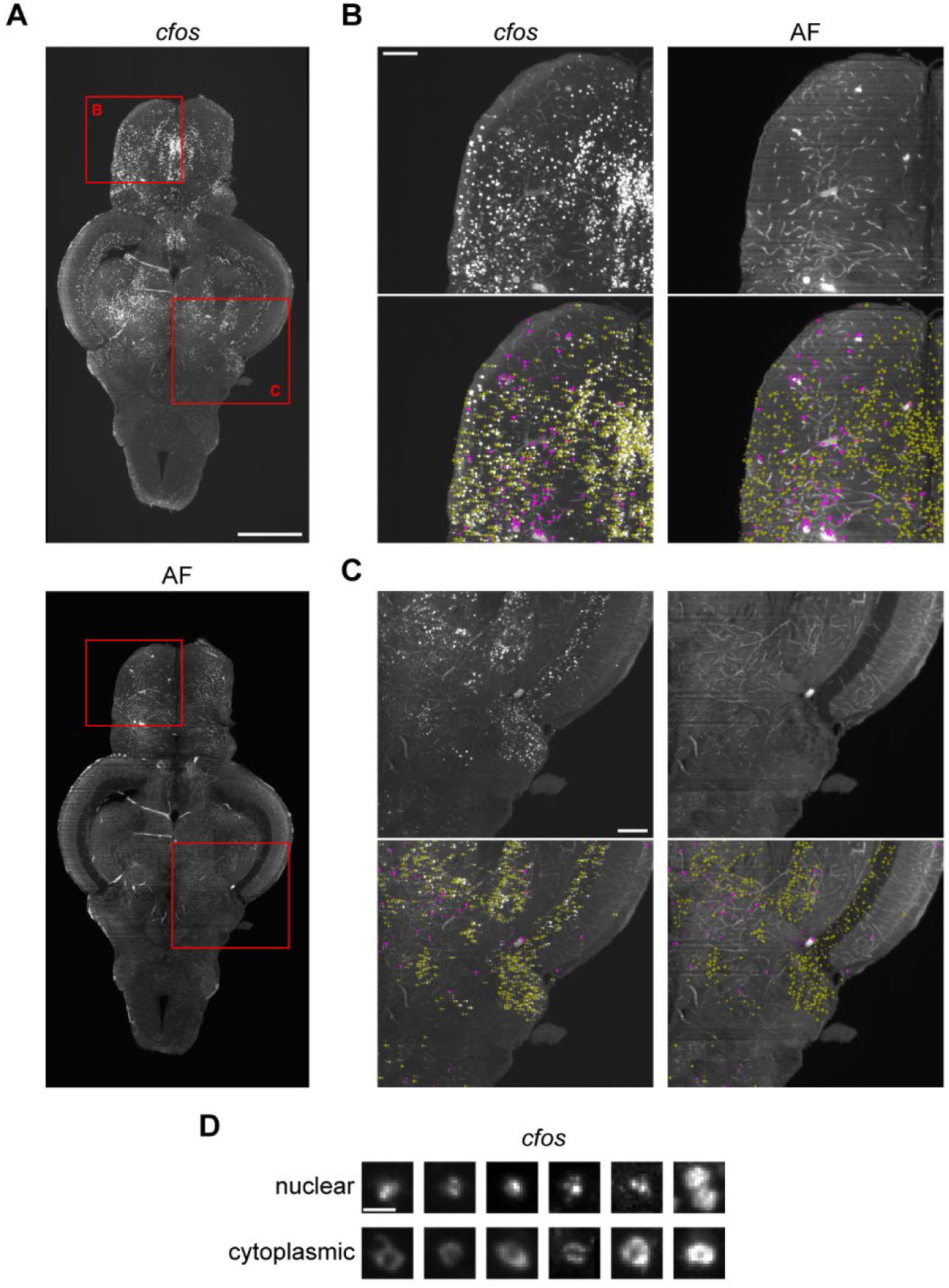
Staining for *cfos* and identifying *cfos* positive cells. A) Adult zebrafish brain stained for *cfos* (top) and the corresponding autofluorescence image (bottom). Scale bar is 0.5 mm. B & C) Zoomed in sections of the brain corresponding to red squares in part A showing *cfos* staining and autofluorescence with labelling of cells (yellow arrows) and non-cells (pink triangles). Scale bars are 0.1 mm. D) Examples of *cfos* staining in the cell nucleus and cytoplasm. Scale bar is 10 μm.

During imaging, we noticed that we had sufficient resolution to differentiate cytoplasmic and nuclear localization of *cfos*. Nuclear staining was characterized by the presence of puncta whereas cytoplasmic staining had a conspicuous dark spot surrounded by more diffuse fluorescence (Figure 2D). This localization of *cfos* is an indication of how long ago the cell was active as the mRNA is first transcribed in the nucleus before being shuttled to the cytoplasm for translation. To capture this distinct cellular localization, we created and trained an artificial neural net on 2,448 examples of nuclear puncta and 1,916 examples of cytoplasmic staining to differentiate these different patterns of *cfos* staining. This network also achieved greater than 95% accuracy.

### Registration to the adult zebrafish brain atlas

The adult zebrafish brain contains over 200 regions, making manual segmentation implausible. To automate parcellation of brains into individual regions, we used ANTs (Avants et al., 2009) to register brains to AZBA using common autofluorescence images. Initially, we attempted to register the autofluorescence image in AZBA directly to individual autofluorescence images, but the results were inconsistent (data not shown). We had more success by first making an average template by registering together 10 autofluorescence images from present study (Figure 3A). The autofluorescence image from AZBA was then successfully registered to this template brain (Figure 3B). A handful of small anomalies arose from this registration process that we manually fixed using ITK-SNAP (Yushkevich et al., 2019). These arose in parts of the image that tend to be highly variable between individuals, such as where mounting occurs at the ventral hypothalamus and the dorsal sac that extends from the dorsal diencephalon. To segment individual brains, we used the transforms from registering the template autofluorescence brain to individual images (Figure 3C). Using inverse transformations from the registration process, we were also able to bring *cfos* images into the space of AZBA (Figure 3D).

**Figure 3.**
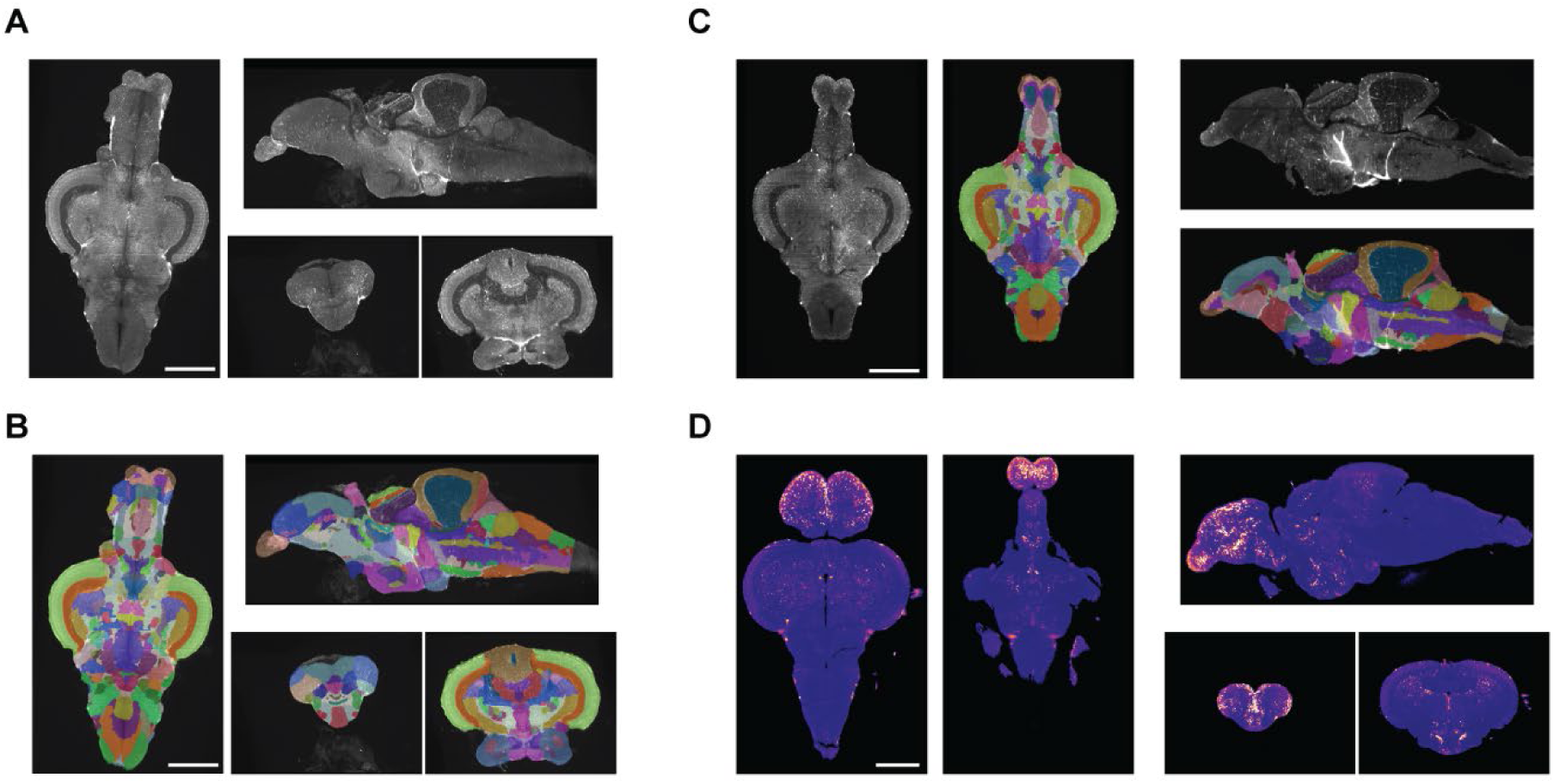
Registration of brain images to AZBA. A) Image of 10 brains registered and averaged. B) Segmentation from AZBA applied to average brain in A. C) Segmentation from AZBA applied to an individual zebrafish brain. D) An individual *cfos* brain brought into the space of AZBA. Scale bars are 0.5 mm.

### Time course for cfos expression

To effectively map whole-brain activity we need to know at what point after behavior *cfos* expression peaks. We exposed fish to a commonly used behavioral task, the novel tank test, and euthanized animals 5, 15, 30, 60, or 120 minutes following the behavior (Figure 4). We also had two control groups: (1) fish that were euthanized immediately after removal from their housing racks, and (2) fish that were brought into the behavioral room and euthanized an hour later, mimicking the habituation to the behavioral room we use for fish that were exposed to the novel tank (i.e., time = 0). A sex × time ANOVA found a large effect of time (P < 0.001, η^2^ = 0.54), a trend towards a small effect of sex (P = 0.07, η^2^ = 0.059), and no interaction (P = 0.46). Using a Dunnet’s t-test to compare all groups to the home tank (HT) control group, we found a large increase in *cfos* cell density at 15 minutes (P = 0.00067, d = 2.07) when *cfos* activity peaked (Figure 4A & B).

**Figure 4.**
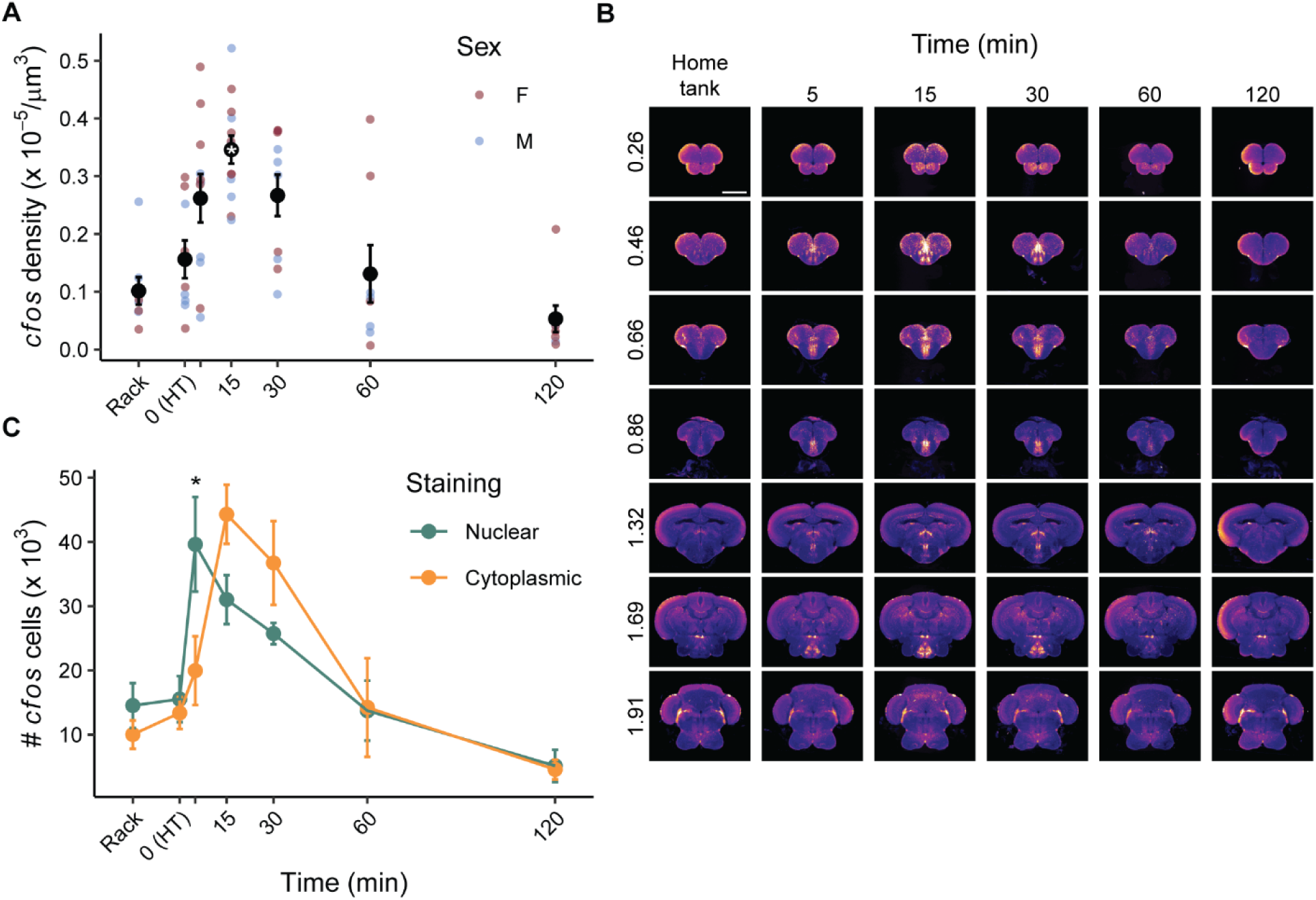
Time course for *cfos* expression following exploration of a novel tank. A) *Cfos* cell density across the entire brain in animals taken off the rack, that remained in their home tank (HT), or a different times after exploration (5, 15, 30, 60, or 120 minutes). * - p < 0.05 compared to the HT group. B) *Cfos* stained brains from each time point were brought into the space of AZBA and averaged and displayed in the coronal plane. The numbers on the left of image are the distance (in mm) from the anterior most portion of the brain. Scale bar is 0.5 mm. C) Number of *cfos* cells classified as nuclear or cytoplasmic at each time point. * - p < 0.05 difference between the number of nuclear and cytoplasmic cells at that time point. Sample sizers were as follows: rack: female: n=4, male: n=4; HT: female: n=5, male: n=4; 5 min: female: n = 6, male: n = 5; 15 min: female: n=7, male: n=6; 30 min: female: n=5, male: n=5; 60 min: female: n=4, male: n=4; 120 min: female: n=5, male: n=3.

We also examined how the proportion of nuclear and cytoplasmic stained cells changed across time (Figure 4C). A cell type × time ANOVA found a large main effect of time (P < 0.001, η^2^ = 0.40) and no overall effect of cell type (P = 0.95). There was also a large interaction between cell type and time (P = 0.0082, η^2^ = 0.13). FDR corrected paired t-tests at each time point found that there were more nuclear than cytoplasmic stained cells at 5 minutes (P = 0.048). This trend switched to more cytoplasmic than nuclear stained cells at 15 and 30 minutes, although the differences at these time points were not statistically significant (P’s = 0.16 & 0.22, respectively).

### Cell types active during the novel tank test

AZBA contains several stains that can be used to identify different cell types across brain regions such as 5-hydroxytryptamine (5-HT), tyrosine hydroxylase (TH), and choline acetyltransferase (ChAT) (Kenney et al., 2021). To determine if exposure to a novel tank results in the activation of regions containing these neuronal cell types, we brought home tank and 15-minute *cfos* brains into the same space as AZBA, averaged the images together, and looked for overlap between the stains in AZBA and elevated *cfos* (Figure 5). For regions expressing 5-HT (Figure 5A), we saw an increase in *cfos* in the paraventricular organ (PVO), intermediate nucleus (IN), and caudal zone of the periventricular hypothalamus (Hc). For TH, which labels dopaminergic and noradrenergic cells, we saw overlap in the ventromedial thalamic nucleus (VM), the posterior part of the parvocellular preoptic nucleus (PPp), paracommissural nucleus (PCN), and Hc (Figure 5B). Finally, for ChAT, we saw overlap in the paraventricular gray zone of the optic tectum (PGZ; Figure 5C). Although we can see overlap at the regional level, our findings are only tentative because the *cfos* and antibody stained images come from separate brains, so we cannot make claims at the cellular level. Nonetheless, this demonstrates how our approach can be used to generate hypotheses about roles different neurotransmitters may play in the underlying a behavior.

**Figure 5.**
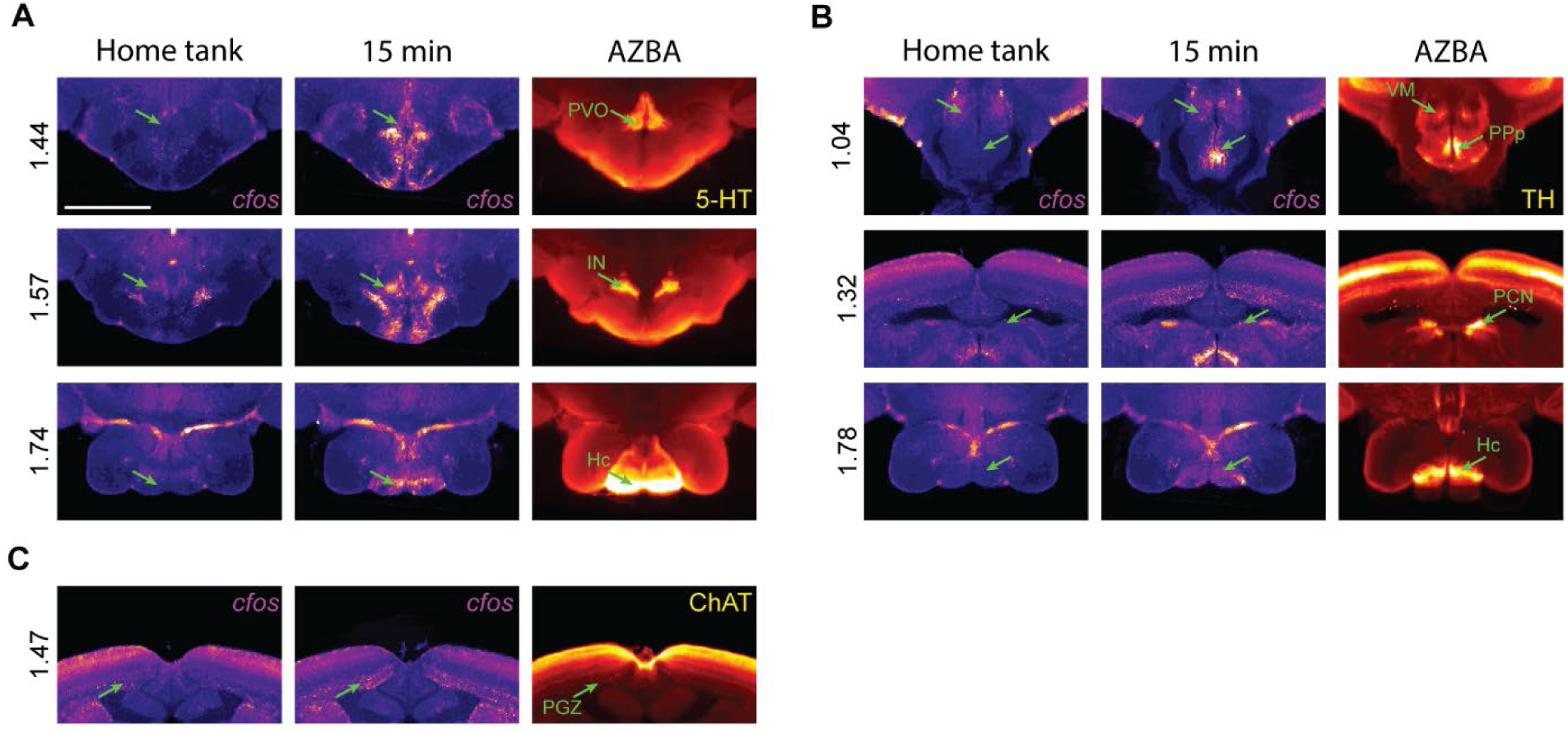
Overlap between *cfos* expression and neurotransmitter-related stains in AZBA. Regional overlap for A) 5-HT, B) TH, and C) ChAT. Scale bar is 0.5mm. Numbers on left are distance from anterior most portion of the brain in mm.

### Brain network analysis

We used functional network analysis to gain insight into the organization of brain activity that underlies exploration of a novel tank (Pinho et al., 2023; Vetere et al., 2017; Wheeler et al., 2013). Using *cfos* counts from the 15-minute time point, we computed the correlated activity between all 143 gray matter regions across animals (Figure 6). To filter the correlation matrix to generate a network, we used efficiency cost optimization where the network density is chosen such that it balances the inclusion of edges to increase global and local efficiency against the putative cost of including additional connections (Fallani et al., 2017). We found a density of 2.5% maximized the efficiency cost optimization quality function (Figure 7A). This resulted in a network with 256 edges and an average degree of 3.6, which is consistent with other functional brain networks generated using different imaging modalities (Fallani et al., 2017). This network also exhibited small world properties: its average shortest path length between nodes was 5.6, which is similar to the average path length of the average from equivalently dense random networks (3.9) with much higher clustering (0.38 versus 0.024). This yielded a small world coefficient greater than 1 (11.0) indicating the expected small world property (Humphries and Gurney, 2008). We also computed degree and eigenvector centrality for each node to uncover brain regions that may play outsized roles in the network (Figure 7C). This uncovered four regions that were in the top 10 for each of these centrality measures: the ventral nucleus of the ventral telencephalon (Vv), the dorsal zone of the ventral telencephalon (Vd-dd), the dorsal most zone of the ventral telencephalon (Vdd), and the anterior part of the parvocellular preoptic nucleus (PPa).

**Figure 6.**
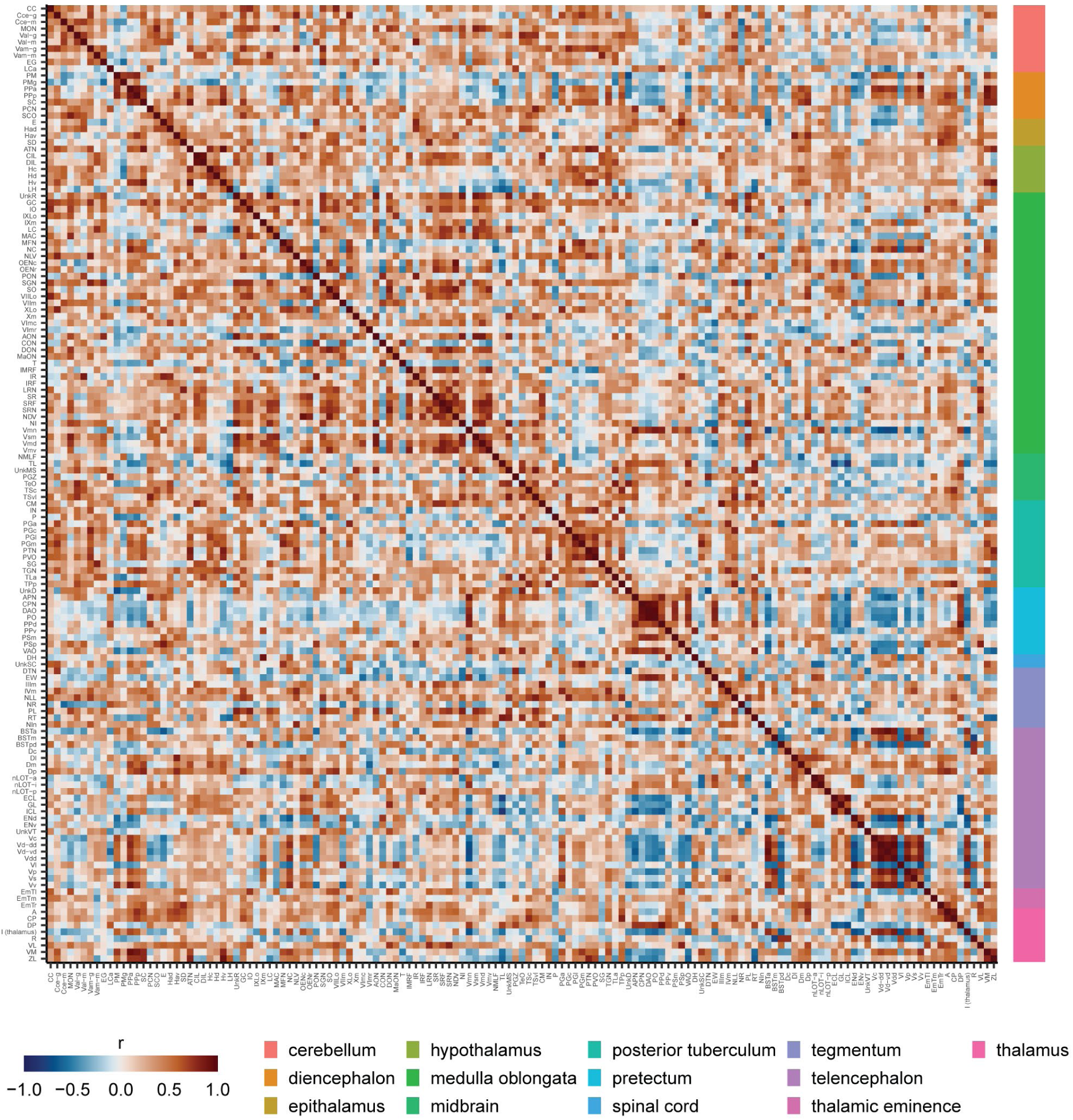
Correlation matrix of *cfos* activity across the zebrafish brain. Entries in the matrix are Pearson correlations between brain regions across animals euthanized 15 minutes after the novel tank test. Regions are organized based on common ontological levels. Regional abbreviations and ontological levels can be found in Table S1.

**Figure 7.**
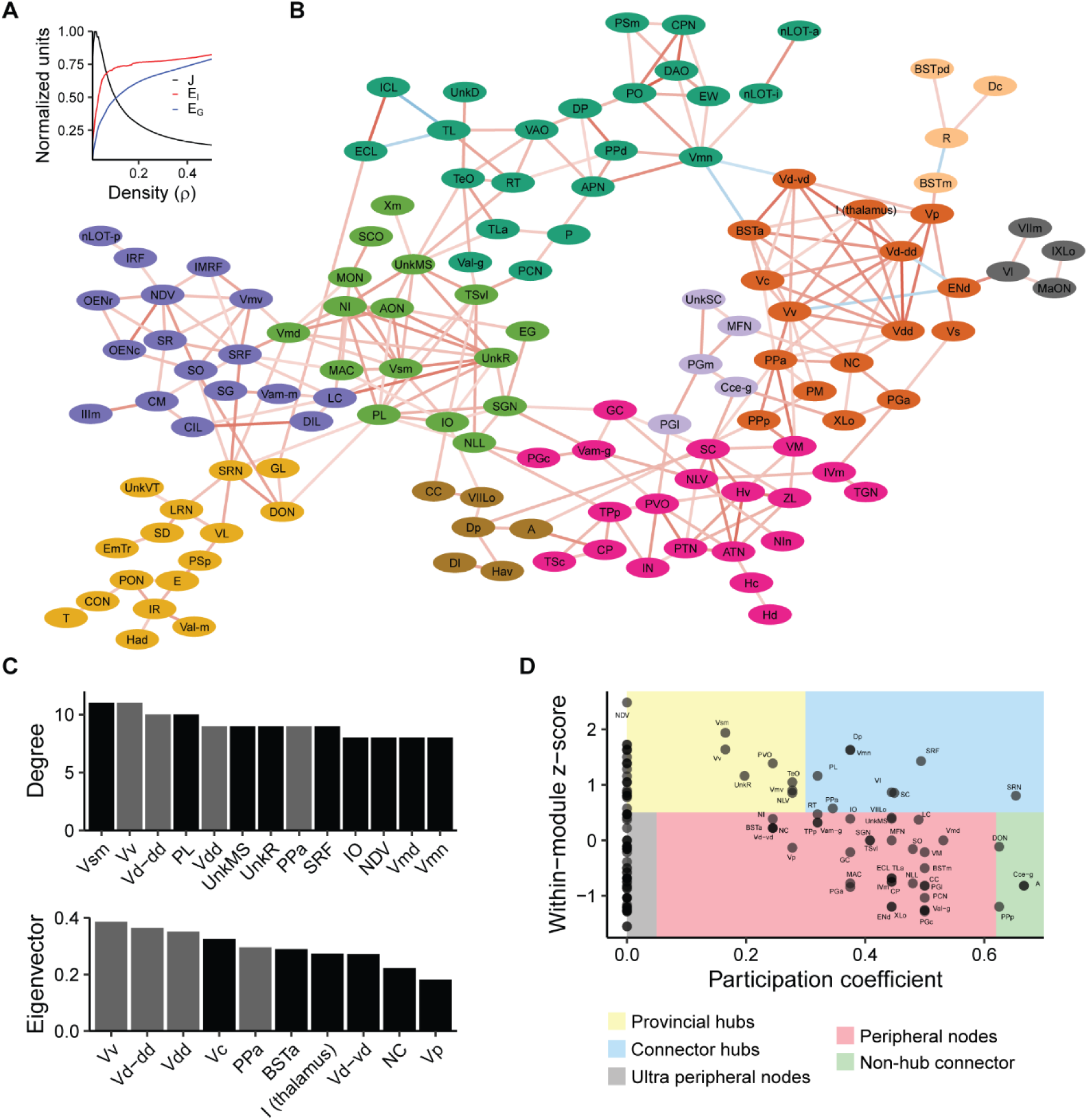
Analysis of the brain network active during the novel tank test. A) Efficiency-cost optimization for different network densities. J: quality function (see methods), E_l_: local efficiency, E_G_: global efficiency. B) Network filtered at a density of 2.5%. Connections between nodes represent suprathreshold correlations from Figure 6. Color of connections represents the strength (darker means higher absolute value) and direction (red: positive, blue: negative) of the correlation. Node colors correspond to communities. Regions not in the giant component are not shown. C) Degree and eigenvector centrality for the top 10 brain regions. Gray bars are those regions that are in the top 10 for both degree and eigenvector centrality. D) Identification of the role that each node plays in the network based on within module degree z-score and participation coefficient.

Next, we used the Louvain algorithm (Blondel et al., 2008) to identify 10 distinct communities in the network (Figure 7B). Using the network and community structure, we categorized the roles that different nodes play in interconnecting different parts of the network (Guimerà and Amaral, 2005): provincial hubs (highly connected within its community, but not between communities), connector hubs (highly connected both within and between communities), peripheral nodes (low connectivity within and between communities), and non-hub connectors (low connectivity within a community, but high between communities).

Interestingly, the PPa, which was identified as important based on centrality measures, arises as a connector hub. The PPa interconnects a module dominated by regions of the ventral telencephalon with other parts of the preoptic area (SC and PPp), thalamus (VM, CP, and ZL) and hypothalamus (ATN, Hv, Hc, and Hd). Thus, our network analysis points to the PPa and ventral telencephalon as likely playing an important role in regulating behavior during exploration of a novel tank.

## Discussion

In the present study, we introduce a pipeline for performing whole-brain activity mapping in adult zebrafish. Our pipeline combines several recently developed tools: a digital brain atlas for adult zebrafish (Kenney et al., 2021), registration using ANTs (Avants et al., 2011), machine learning tools for automated cell detection (Tyson et al., 2021), tissue clearing (Renier et al., 2014), light-sheet microscopy (Reynaud et al., 2014), and *in situ* HCR (Choi et al., 2018) for detecting *cfos*. Importantly, all the computational tools are open access and free to use.

Furthermore, to aid in the implementation of this pipeline, we have included a bench protocol (Supplemental file 1). The primary stumbling blocks for implementing this pipeline are likely to be access to a light-sheet microscope for whole-brain imaging and sufficient computational power for training and applying the registration and CellFinder machine learning algorithms. The former issue is partly mitigated by the increased availability of light-sheet microscopes, particularly in core facilities. Access to computational resources can be addressed by using tools like Google Colaboratory (Bisong, 2019) or high performance computing facilities available at many institutions.

### Cfos to capture whole-brain activity

We captured neural activity using *in situ* HCR to detect *cfos* mRNA. We chose this approach for several reasons: (1) there are a paucity of antibodies for detecting cfos protein in zebrafish, none of which are known to work in whole-mount tissue-cleared samples, (2) *in situ* HCR probes are small (∼150 bp), which easily penetrates chunks of intact tissue like the adult zebrafish brain, and (3) *cfos* is one of the most widely used markers of neural activity due to autoinhibition of transcription that results in low background, high signal-to-noise, and good temporal resolution (Chung, 2015; Lucibello et al., 1989). The findings in the present study further support these rationales: we saw even penetration of *cfos* staining throughout the brain (Figures 2 and 4B) and the levels of background *cfos* staining were low, with an approximately 3.5 fold increase in *cfos* density 15 minutes following behavior compared to quiescent animals removed directly from their housing racks (Figure 4A). The increase in *cfos* was also tightly coupled to the behavior, peaking 15 minutes after exposure to the novel tank before decreasing to baseline levels by 60 minutes. Interestingly, if we look at only cells that have nuclear staining, we see the increase begins as soon as 5 minutes after behavior. The higher *cfos* density at 15 minutes is likely due to the opportunity for increased transcription which would be expected to create a brighter signal resulting a larger number of detectable cells. The time to maximal *cfos* we observed is faster than is seen in rodents, where it is often found to peak at 30 minutes post-stimulation (Ding et al., 1994; Guzowski et al., 2001; Kovács, 1998; Zangenehpour and Chaudhuri, 2002). The reasons for this time difference between zebrafish and rodents is unclear. Nonetheless, it emphasizes the importance of performing time course analysis when establishing new methods for brain mapping in different species.

Other markers of neural activity have gained traction in recent years in zebrafish, such as the phosphorylated forms of ribosomal protein S6 (pS6) and extracellular regulated kinase 1/2 (pERK1/2). Our data suggests that *cfos* as an activity marker compares favorably to these options. For pS6, the signal-to-noise ratio is comparable to what we see for *cfos*, with an approximately 2-4 fold increase over baseline both *in vivo* in zebrafish (Butler et al., 2018; Parada et al., 2024; Scaia et al., 2022) and *in vitro* neuronal cell culture (Kenney et al., 2015). However, the time course of elevated pS6 is notably slower, taking an hour or more to peak (Kenney et al., 2015; Parada et al., 2024) compared to 15 minutes for *cfos* (Figure 4). In contrast, pERK1/2 activity peaks quickly, within 2-5 minutes, but the signal-to-noise ratio is ∼0.5-1, considerably lower than *cfos* (Randlett et al., 2015; Venincasa et al., 2021). This low signal-to-noise ratio likely arises from higher background levels of pERK due to the wide variety of cellular processes that it regulates (Cargnello and Roux, 2011). Thus, the best choice of stain depends on the behavioral paradigm. Large, rapid responses to brief behavioral stimuli are best captured by pERK. However, more subtle responses may be missed due to the low signal-to-noise ratio. S6 phosphorylation excels at capturing long lasting steady-state neural activity, as suggested by Maruska et al (2020) and would excel for behaviors lasting 30 minutes or more.

C*fos* represents a solid middle ground that is ideal for capturing neural activity from behaviors lasting on the order of 5-10 minutes, like the novel tank test used in the present study.

### Registration to AZBA to identify cell types

We were able to successfully register our brains to AZBA using ANTs (Avants et al., 2009). To do so, we first used ANTs to make an average template from our images by registering 9 brains to a single brain and averaging them together. The autofluorescence image in ABZA was then registered to this average template, yielding good results (Figure 3). We chose this method because we found that registering the autofluorescence image from AZBA to individual brains gave inconsistent results. This is likely because the autofluorescence image in AZBA is also an average of many brains (Kenney et al., 2021). We chose ANTs because the non-linear symmetric diffeomorphic image registration it employs has been consistently found to be one of the best algorithms for 3D image registration (Klein et al., 2009; Murphy et al., 2011). The tool is also well documented and straightforward to use. Finally, ANTs has recently grown in popularity for image registration in larval zebrafish (Marquart et al., 2017; Shainer et al., 2023), which provided a starting point for identifying the best parameters for registration in our samples.

Following registration to AZBA, we were able to identify potential neuronal cell types relevant to the novel tank test (Figure 5). We found that several regions containing high levels of 5-HT were active during behavior, such as the PVO, IN, and Hc. Consistent with this, several papers have implicated 5-HT as contributing to exploration of a novel tank using pharmacological approaches (Beigloo et al., 2024; Maximino et al., 2013; Nowicki et al., 2014; Wong et al., 2010). Similarly, there was overlap in *cfos* activity in several regions that express tyrosine hydroxylase (VM, PPp, PCN, and Hc), implicating these populations of dopaminergic or noradrenergic neurons in novel tank behavior (Kacprzak et al., 2017; Nabinger et al., 2023). Of the *cfos* positive cells that overlap with TH, our network analysis suggests that the PPp may be of particular importance in regulating exploratory behavior, as it is one of the few non-hub connectors (Figure 7D). The PPp also has a direct connection to the PPa region, which ranks highly in both eigenvector and degree centrality (Figure 7B), and connects to the thalamic VM region, another area high in TH expression. This suggests that the PPp and VM may may act in concert to mediate the effects of the dopaminergic system on exploration. However, one important caveat to these interpretations is that we are comparing averaged *cfos* images to averaged neurotransmitter-related stains in AZBA, and thus we cannot definitively identify the specific cell types that are active. This would require co-staining of brains with both *cfos* and various neuronal cell-type markers to determine if the activity of these specific cell types changes.

### Novel tank functional connectome

Using our whole brain mapping data, we generated the first novel tank functional connectome. The novel tank test is one of the most widely used behavioral tests in adult zebrafish, often used to study exploratory and anxiety-related behaviors (Blaser et al., 2010; Kalueff et al., 2013; Luca and Gerlai, 2012; Rajput et al., 2022; Spence et al., 2006). Our functional network analysis identifies several key regions that are engaged during exploration of a novel tank for the first time (Figures 6 and 7). In particular, the medial portion of the ventral telencephalon stands out, where several subregions (the Vv, Vd-dd, Vc, Vd-vd, and Vp) rank highly on at least one measure of centrality (Figure 7C). These regions are also highly interconnected, a fact that is clear from both the correlation matrix (Figure 6) and the community they form in the network (dark orange in Figure 7B). Based on molecular markers, these regions of the ventral telencephalon are thought to correspond to the mammalian subpallial amygdala (i.e., the central and medial amygdala) and basal ganglia (Mueller, 2022; Porter and Mueller, 2020). In mammals, these brain regions have been found to be important for a wide range of behaviors, from defensive, anxiety-related, and social behaviors to motor control (Fadok et al., 2018; Grillner and Robertson, 2016; Raam and Hong, 2021). Our findings that the ventral telencephalon appears to be engaged during the novel tank test is reasonable given that novelty and exploration would be expected to engage circuits involved in decision making, emotional regulation, and muscle coordination.

In examining how the regions of the ventral telencephalon interact with the rest of the brain, a few interesting trends emerge. Notably, the interaction of ventral telencephalic regions with many other communities is anti-correlated (i.e., the dark green, light orange, and grey communities in Figure 7B). This suggests the presence of strong inhibitory connections between the medial ventral telencephalon and other parts of the brain. Consistent with this interpretation, the ventral telencephalon has been found to contain a substantial number of inhibitory GABAergic neurons (Porter and Mueller, 2020). Our network analysis suggests that these inhibitory connections are most likely present between the ventral telencephalon and the Vmn (mesencephalic nucleus of the trigeminal nerve), End (entopeduncular nucleus in the lateral portion of the ventral telencephalon), and from the BSTm (bed nucleus of the stria terminalis, medial portion in the dorsal telencephalon) to R (rostrolateral nucleus in the thalamus). However, given that our findings are correlational in nature, techniques like tract tracing and direct manipulation would be needed to confirm these interactions.

Our network analysis also identified the PPa as a region of high importance. The PPa was high in both eigenvector and degree centrality (Figure 7C) and was one of the few connector hub nodes (Figure 7D). In examining its place in the network (Figure 7B), the PPa interconnects with several regions of the ventral telencephalon and, working in concert with the PPp, mediates their interactions with parts of the network that contain several thalamic and hypothalamic regions (magenta cluster in Figure 7B). To our knowledge, the correspondence between the PPa and PPp in teleosts and tetrapods has not been determined. Based on the expression of neuropeptides like oxytocin and arginine vasopressin, parts of the PPp are thought to be equivalent to the supraoptic nucleus in mammals (Herget et al., 2014). In larval zebrafish, the preoptic area has recently been implicated in behaviors such as navigation, thermoregulation, and stress reactivity (Corradi et al., 2022; Palieri et al., 2024). However, the preoptic area in larval zebrafish cannot be differentiated into subregions like the PPa and PPp due to a lack of cytoarchitectural boundaries (Herget et al., 2014). This makes it unclear as to what specific regions in the adult would subsume the functions identified in larval animals. Future work should determine the role that these different subregions might play in different aspects of exploration and anxiety-like behavior in adult zebrafish.

### Summary

The present study provides an open-source framework for performing whole-brain mapping in adult zebrafish. This work also yielded the first description of brain activity that underlies the novel tank test, suggesting the medial ventral telencephalon may play an important role in one of the most widely used behavioral tasks in adult zebrafish. Taken together, we anticipate that our pipeline will help generate insights into the principles of brain function by enhancing the utility of adult zebrafish as a model organism.

## Methods

### Animals

#### Zebrafish

Subjects were 8–10 month old zebrafish of the TU strain from both sexes. Fish were bred and raised at Wayne State University and within two generations of animals obtained from the Zebrafish International Resource Center (ZIRC, catalog ID: ZL84) at the University of Oregon. Fish were maintained in high-density racks under standard conditions: water temperature of 27.5 ± 0.5 °C, salinity of 500 ± 10 µS, and pH of 7.4 ± 0.2. Lighting followed a 14:10 light:dark cycle, with lights on at 8:00 AM. Fish were fed twice daily with dry feed (Gemma 300, Skretting, Westbrook, ME, USA) in the morning and brine shrimp (Artemia salina, Brine Shrimp Direct, Ogden, UT, USA) in the afternoon.

Sex determination was based on secondary sex characteristics such as shape, color, and the presence of pectoral fin tubercles (McMillan et al., 2015). Confirmation was conducted post-experimentation by euthanizing the animals and observing the presence or absence of eggs. All experimental procedures were conducted under the ethical approval of the Wayne State University Institutional Animal Care and Use Committee (Protocol ID: 21-02-3238).

#### Behavioral stimuli and tissue collection

The novel tank test was used as the behavioral stimulus, using tanks that were distinct from housing tanks. Behavioral tanks were open top five-sided (15 × 15 × 15 cm) and made from frosted acrylic (TAP Plastics, Stockton, CA, USA). Each tank was filled to a height of 12 cm with 2.5 L of fish facility water and housed within a white corrugated plastic enclosure to minimize external disturbances and diffuse light.

One week before the novel tank test, animals were housed in 2-liter tanks divided into two chambers with transparent dividers. Male and female pairs were kept in each chamber to enable identification of individuals without social isolation or tagging. A day prior to the experiment, animals were acclimatized to the behavior room for one hour before being placed back on the housing racks. On the day of the experiment, animals were removed from the housing rack and allowed to acclimate in the behavioral room for one hour. After acclimation, animals were individually transferred to a novel tank and allowed to explore the tank for 6 minutes. Water was replaced between animals. After six minutes, fish were removed and placed back in their home tank for a designated periods of time (5, 15, 30, 60, or 120 minutes) prior to euthanization. A subset of animals was euthanized one hour after acclimation to the room (home tank control) and another set of animals were euthanized immediately after removal from the housing racks (rack control).

Animals were euthanized by immersion in ice cold water for 5 minutes to induce anesthesia and then decapitated using a sharp blade. Heads were then washed in ice-cold phosphate buffered saline (PBS) for 60 seconds to allow for blood drainage, and then fixed in 4% paraformaldehyde in PBS overnight. Brains were then dissected in ice cold PBS and subject to iDISCO+ and *in situ* HCR.

### Histology

#### Tissue pre-treatment

We adapted the iDISCO+ protocol (Renier et al., 2016) for zebrafish brain tissue staining. Following dissection, brain samples were washed for 30 minutes, three times, in PBS at room temperature. This was followed by dehydration using a methanol concentration gradient (20, 40, 60, 80, and 100%) for 30 min each. Samples were further washed in 100% methanol, chilled on ice, and then incubated in chilled 5% hydrogen peroxide in methanol overnight at 4°C. The next day, the samples were rehydrated through a reverse methanol series (80%, 60%, 40%, 20%) at room temperature, followed by a 1 h PBS wash, two 1 h PBS-T washes (1x PBS, 0.1% Tween 20), and a 3 h PBS-T wash. Samples were then equilibrated overnight in 5× SSCT (sodium chloride sodium citrate/0.1% Tween-20) buffer.

#### In-situ HCR

We modified the original HCR method described by Choi and colleagues (2018) and informed by the work of Kumar et al (2021). Samples were first prepared by acetylation in 0.25% v/v acetic anhydride solution in ultrapure water for 30 min. Samples were then washed in ultrapure water three times for 5 mins and then equilibrated in probe hybridization buffer (30% formamide, 5x SSC, 9 mM citric acid, 0.1% Tween-20, 50 µg/mL heparin, 1x Denhardt’s solution, 10% Dextran sulfate) for 15 min at room temperature. Samples were then incubated in probe hybridization buffer for 1h at 37 °C with shaking and then incubated with 1 µM of *cfos* probes in hybridization buffer at 37 °C with shaking for 48-60 hours. Samples were then washed with probe wash buffer (30% formamide, 5x SSCT, 9 mM citric acid, 50 µg/mL heparin) three times at 37°C, then twice with 5× SSCT for 1 h each with shaking. The tissue was then equilibrated in amplification buffer (5x SSC, 0.1% Tween-20, 10% Dextran sulfate) at room temperature for 1h with shaking. Alexa647 labeled hairpins (B1) were prepared by heating to 95 °C for 90 seconds prior to cooling at room temperature in the dark. We diluted 7.5 pmol of each hairpin into 125 µL of amplification buffer for each sample. Samples were incubated for 48-60 hours in the dark at room temperature. Finally, tissue was washed in 5× SSCT, 5 times for 1 hour each before being washed overnight in 5X SSCT.

#### Tissue clearing

Samples were first dehydrated in a series of methanol-water mixtures (20%, 40%, 60%, 80%, 100%) at room temperature for 1 hour each and then left in 100% methanol overnight. The next day, samples were incubated at room temperature in a mixture of 66% dichloromethane and 33% methanol for 3 hours followed by two 15-minute washes in dichloromethane. After removing the dichloromethane, samples were incubated and stored in dibenzyl ether at room temperature for at least 24 hours until imaging.

#### Brain imaging and processing

Cleared samples were imaged on a LaVision BioTec UltraMicroscope II (Miltenyi Biotec, Auburn, CA) using Imspector software for image acquisition. The microscope setup included a 4.2 Megapixel sCMOS camera and a 2x objective lens with a dipping cap with spherical aberration correction. Images were taken at a magnification of 6.4x. Samples were mounted on the sample holder using an ultraviolet cured resin (NOA 61, Norland Products, Jamesburg, NJ) with a refractive index (1.56) that matched DBE. Imaging was conducted from the right laser sheet with a 4 μm step size using dynamic horizontal focus. Both 480 nm autofluorescence and 640 nm signal channels were used. The imaging settings used were: 90% laser power, 200-ms exposure time, 50% sheet width, sheet numerical aperture of **<u>XX</u>**. Acquired images were stitched using Terastitcher (Bria and Iannello, 2012).

### Computational analysis

#### Automated cell detection

For the automated detection and quantification of *cfos* positive cells, we utilized the Python-based software, CellFinder (Tyson et al., 2021). It comprises two steps: cell candidate detection and cell classification. The initial step of cell detection identifies cell-like objects in the image. We optimized parameters to capture as many cell-like objects in our images as possible. Running from the Linux terminal, we used the following command for cell detection:

~~~
cellfinder -s path/to/folder/signal/channel/cfos -b/path/to/folder/background/channel/AF -o path/to/output1 -v 3.990 0.943 0.943 --orientation sal --no-register --no-classification -- soma-diameter 5 --threshold 3 --ball-xy-size 2 --ball-z-size 7 --ball-overlap-fraction 0.77 --log-sigma-size 0.1 --save-csv --batch-size 64 --epochs 100
~~~

After detecting cell candidates, a customized python script was used to remove cell candidates that were within 9 μm of one another.

Napari was utilized for visualization and labelling. We manually annotated 10,597 cells and 7,303 non-cells across five brains for training the artificial neural network. Cellfinder was trained using the following command:

~~~
Cellfinder_train -y path/to/brain1_labels.yml path/to/brain2_labels.yml path/to/brain3_labels.yml path/to/brain4_labels.yml path/to/brain5_labels.yml -o/trained_network --batch-size 64 --epochs 100 --no-save-checkpoints -- save-progress
~~~

The trained network achieved 96.1% accuracy. Finally, the trained network was applied to all the experimental brains to classify the detected cell candidates into cells and non-cells. This was achieved by utilizing the following command:

~~~
cellfinder -s /path/to/folder/signal/channel/cfos/ -b/path/to/folder/background/channel/AF/ -o path/to/output -v 3.990 0.943 0.943 --orientation sal --no-register --no-detection --soma- diameter 5 --threshold 3 --ball-xy-size 2 --ball-z-size 7 --ball-overlap-fraction 0.77 --log-sigma-size 0.1 --save-csv --trained-model/trained_network/model.h5
~~~

#### Differentiating nuclear and cytoplasmic stained cells

To differentiate between cytoplasmic and nuclear puncta, we developed a convolutional neural network (CNN) built in Python using the TensorFlow library. The architecture of the CNN is outlined in Table S2. *Cfos* Images from 10 brains were labelled, totaling 2,448 puncta and 1,916 cytoplasmic labels. A training dataset was created by isolating 11ˣ11ˣ11 pixel cubes centered around each of the labeled cells. The dataset was split 80/20 into a training set and a testing set. The input data was augmented through a series of horizontal and vertical flips, 90° rotations, and 2-pixel horizontal translations to create a total training dataset of 13,706 puncta and 10,724 cytoplasmic labels. No data augmentation was performed on the testing set. The model was trained using an NVIDIA GeForce 3090 GPU for 500 epochs. The batch size was 32, the weight decay rate was 0.0005, and the learning rate was 0.0001. The model achieved an accuracy of 95.3% on the testing set.

#### Brain registration

Image registration was performed using ANTs (Avants et al., 2009). For the non-linear diffeomorphic step, four parameters were optimized: cross-correlation, gradient step, update field variance in voxel space, and total field variance in voxel space to achieve the best alignment. Using the optimized parameters, brain registration was carried out in two steps: first, an average brain template was created, and second, AZBA was registered to this average template.

Before registration, images were downsampled to 4 μm isotropic using brainreg from the BrainGlobe suite of tools (Tyson et al., 2021):

~~~
brainreg /path/to/raw/data /path/to/output/directory -v 3.990 0.943 0.943 --orientation sal --atlas azba_zfish_4um –debug
~~~

The average template was generated using 10 autofluorescence images. Initially, nine autofluorescence images were individually brought into the space a single image (template) using the following ANTs command:

~~~
antsRegistration --dimensionality 3 --float 1 -o [${AF_sample_1_for_avg_},${ AF_sample_1_for_avg-warped}] -- interpolation WelchWindowedSinc -u 0 -r [${AF_template.nii},${AF_sample_1.nii},1] -t Rigid[0.1] -m MI[${AF_template.nii},${AF_sample_1.nii},1,32,Regular,0.25] -c [200 × 200 × 200 × 0,1e-8,10] --shrink-factors 12x8x4x2 --smoothing-sigmas 4x3x2x1vox -t Affine[0.1] -m MI[${AF_template.nii},${AF_sample_1.nii}, 1,32,Regular,0.25] -c [200 × 200 × 200 × 0,1e-8,10] --shrink-factors 12x8x4x2 --smoothing-sigmas 4x3x2x1vox -t SyN[0.3,4,0] -m CC[${AF_template.nii},${AF_sample_1.nii}, 1,3] -c [200 × 200 × 200 × 200, 1e-6,10] --shrink-factors 12x8x4x2 --smoothing-sigmas 4x3x2x1vox --verbose 1
~~~

These outputs were then used to create an average image using the ‘AverageImages’ command in ANTs. Next, the autofluorescence image from AZBA was registered to the average template using the following command:

~~~
antsRegistration --dimensionality 3 --float 1 -o [${AZBA_to_avg_temp_},${AZBA_to_avg_temp-warped}] --interpolation WelchWindowedSinc -u 0 -r [${avg_template.nii.gz},${AZBA/20180628_AF_average.nii.gz},1] -t Rigid[0.1] -m MI[${avg_template.nii.gz},${AZBA/20180628_AF_average.nii.gz},1,32,Regu lar,0.25] -c [200 × 200 × 200 × 0,1e-8,10] --shrink-factors 12x8x4x2 --smoothing-sigmas 4x3x2x1vox -t Affine[0.1] -m MI[${avg_template.nii.gz},${AZBA/20180628_AF_average.nii.gz}, 1,32,Regular,0.25] -c [200 × 200 × 200 × 0,1e-8,10] --shrink-factors 12x8x4x2 --smoothing-sigmas 4x3x2x1vox -t SyN[0.3,4,0] -m CC[${avg_template.nii.gz},${AZBA/20180628_AF_average.nii.gz}, 1,3] -c [200 × 200 × 200 × 200, 1e-6,10] --shrink-factors 12x8x4x2 -- smoothing-sigmas 4x3x2x1vox --verbose 1
~~~

To bring the segmentation from AZBA into the space of the template we used the following command:

~~~
antsApplyTransforms -d 3 --float -n NearestNeighbor -i /AZBA/2021-08-22_AZBA_segmentation.nii.gz -r avg_template.nii.gz -o AZBA_to_avg_temp_transformed.nii.gz -t AZBA_to_avg_temp_1Warp.nii.gz - t AZBA_to_avg_temp_0GenericAffine.mat
~~~

Finally, the newly generated average template image was used as a reference image and was registered onto individual autofluorescence images:

~~~
antsRegistration --dimensionality 3 --float 1 -o [${AF_sample_},${AF_sample-warped}] --interpolation WelchWindowedSinc -u 0 -r [${AF_sample.nii},${avg_template.nii.gz },1] -t Rigid[0.1] -m MI[${AF_sample.nii},${ avg_template.nii.gz },1,32,Regular,0.25] -c [200 × 200 × 200 × 0,1e-8,10] --shrink-factors 12x8x4x2 --smoothing-sigmas 4x3x2x1vox -t Affine[0.1] -m MI[${AF_sample.nii},${ avg_template.nii.gz },1,32,Regular,0.25] -c [200 × 200 × 200 × 0,1e-8,10] --shrink-factors 12x8x4x2 --smoothing-sigmas 4x3x2x1vox -t SyN[0.3,4,0] -m CC[${AF_sample.nii},${ avg_template.nii.gz },1,3] -c [200 × 200 × 200 × 200, 1e-6,10] --shrink-factors 12x8x4x2 -- smoothing-sigmas 4x3x2x1vox --verbose 1
~~~

Finally, segmentation of individual brains was done using the same command as above but applied to the segmentation file as the floating image.

#### Cfos cell counts and network analysis

R (version 4.1.1) was used for network analysis and to integrate the output from Cellfinder with the brain segmentation using the RNifti package (Clayden et al., 2021) to read in the segmentation files. The number *cfos* positive cells in each brain were summed excluding white matter and clear labelled regions yielding 143 gray matter regions for analysis.

Network analysis was performed using the igraph (version 2.0.2) package (Csardi and Nepusz, 2006). The network was generated by treating the correlation matrix (Figure 6) as an adjacency matrix. For thresholding we chose the network density using efficiency cost optimization to maximize the quality function (Fallani et al., 2017):

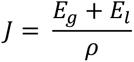

Where *E*_g_ is the global efficiency, *E*_l_ is the average of the local efficiency, and *ρ* is the network density. For the calculations of global and local efficiency we used a binarized network based on the absolute value of the correlations.

For identifying node roles, we calculated the within module degree z-score:

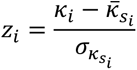

Where *κ_i_* is the number of connections between node *i* and other nodes in the same community and *κ̅_si_* is the average of over all nodes in a community; *σ*_*κs_i_*_ is the standard deviation of the number of connections in a community. We also calculated the participation coefficient:

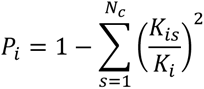

Where *N_c_* is the number of communities, *K*_*is*_ is the number of connections between node *i* and all other nodes in community s, and *K*_*i*_ is the degree of node *i*. The definitions of the above equations and the boundaries for the different types of nodes were based on the Guimerà and Amaral (2005).

The small worldness parameter was calculated as described in (Humphries and Gurney, 2008):

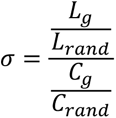

Where *L*_*g*_ is the average shortest path length between all nodes of the network, *L_rand_* is the average shortest path length between all nodes in an equivalent random network, *C*_*g*_ is the clustering coefficient of the network, and *C_rand_* is the clustering coefficient of an equivalent random network. For random network parameters, we took the average from 1,000 instances of Edros-Renyi random networks (Erdös and Rényi, 2011) with an equivalent number of nodes and edges as the target network.

### Statistical analysis

Statistical analysis was done using R. Data were analyzed using 2 × 2 ANOVAs as indicated in the results. For the overall time course *cfos* data, Dunnet’s t-tests were used to compare all other groups to the home tank control group (time = 0). False discovery rate (FDR; Benjamini and Hochberg, 1995) corrected paired t-tests at each time point were used for cytoplasmic versus nuclear data.

## Supporting information

Supplemental File 1: Bench protocol

## Data and code availability

Data and code are available at github: https://github.com/KenneyLab/RajputEtAl_2024_Whole_brain_mapping

## Acknowledgments and Funding

We thank Jacob Hudock and Dinh Luong for excellent care of the zebrafish and facility maintenance. This work was supported by the National Institutes for General Medical Sciences (R35GM142566, JWK) and the Richard Barber Interdisciplinary Research Program (KKF, JWK).

