## Supplemental tables for "Whole-brain mapping in adult zebrafish and identification of a novel tank test functional connectome"

Table S1. List of abbreviations for grey matter regions

| Abbreviation | Name | Ontological level |
| --- | --- | --- |
| A | anterior thalamic nucleus | thalamus |
| AON | anterior octaval nucleus | medulla oblongata |
| APN | accessory pretectal nucleus | pretectum |
| ATN | anterior tuberal nucleus | hypothalamus |
| BSTa | bed nucleus of the stria terminalis, anterior division | telencephalon |
| BSTm | bed nucleus of the stria terminalis, medial division | telencephalon |
| BSTpd | bed nucleus of the stria terminalis, posterior division | telencephalon |
| CC | cerebellar crest | cerebellum |
| Cce-g | cerebellar corpus, granular layer | cerebellum |
| Cce-m | cerebellar corpus, molecular layer | cerebellum |
| CIL | central nucleus of the inferior lobe | hypothalamus |
| CM | mammillary body | posterior tuberculum |
| CON | caudal octavolateralis nucleus | medulla oblongata |
| CP | central posterior thalamic nucleus | thalamus |
| CPN | central pretectal nucleus | pretectum |
| DAO | dorsal accessory optic nucleus | pretectum |
| Dc | central zone of dorsal telencephalon area | telencephalon |
| DH | dorsal horn | spinal cord |
| DIL | diffuse nucleus of the inferior lobe | hypothalamus |
| Dl | lateral zone of the dorsal telencephalon | telencephalon |
| Dm | medial zone of dorsal telencephalon | telencephalon |
| DON | descending octaval nucleus | medulla oblongata |
| Dp | posterior zone of dorsal telencephalon area | telencephalon |
| DP | dorsal posterior thalamic nucleus | thalamus |
| DTN | dorsal tegmental nucleus | tegmentum |
| E | epiphysis | epithalamus |
| ECL | external cellular layer of olfactory bulb | telencephalon |
| EG | granular eminence | cerebellum |
| EmTl | lateral thalamic eminence | thalamic eminence |
| EmTm | medial thalamic eminence | thalamic eminence |
| EmTr | rostral thalamic eminence | thalamic eminence |
| ENd | entopeduncular nucleus, dorsal part | telencephalon |
| ENv | entopeduncular nucleus, ventral part | telencephalon |
| EW | Edinger-Westphal nucleus | tegmentum |
| GC | central gray | medulla oblongata |
| GL | glomerular layer of olfactory bulb | telencephalon |
| Had | dorsal habenular nucleus | epithalamus |
| Hav | ventral habenular nucleus | epithalamus |
| Hc | caudal zone of periventricular hypothalamus | hypothalamus |
| Hd | dorsal zone of periventricular hypothalamus | hypothalamus |
| Hv | ventral zone of periventricular hypothalamus | hypothalamus |
| I (thalamus) | Intermediate thalamic nucleus | thalamus |
| IAF | inner arcuate fibers |  |
| ICL | internal cellular layer of olfactory bulb | telencephalon |
| IMRF | intermediate reticular formation | medulla oblongata |
| IN | Intermediate nucleus | posterior tuberculum |
| IO | inferior olive | medulla oblongata |
| IR | inferior raphe | medulla oblongata |
| IRF | inferior reticular formation | medulla oblongata |
| LC | locus coeruleus | medulla oblongata |
| LCa | caudal lobe of cerebellum | cerebellum |
| LH | lateral hypothalamic nucleus | hypothalamus |
| LRN | lateral reticular nucleus | medulla oblongata |
| MAC | Mauthner cell | medulla oblongata |
| MaON | magnocellular octaval nucleus | medulla oblongata |
| MFN | medial funicular nucleus | medulla oblongata |
| MON | medial octavolateralis nucleus | cerebellum |
| NC | commissural nucleus of Cajal | medulla oblongata |
| NDV | nucleus of the descending trigeminal root | medulla oblongata |
| NI | isthmic nucleus | medulla oblongata |
| NIn | interpeduncular nucleus | tegmentum |
| NLL | nucleus of the lateral lemniscus | tegmentum |
| nLOT-a | nucleus of the lateral olfactory tract, anterior part | telencephalon |
| nLOT-i | nucleus of the lateral olfactory tract, intermediate part | telencephalon |
| nLOT-p | nucleus of the lateral olfactory tract, posterior part | telencephalon |
| NLV | nucleus lateralis valvulae | medulla oblongata |
| NMLF | nucleus of the medial longitudinal fascicle | midbrain |
| NR | red nucleus | tegmentum |
| OENc | octavolateralis efferent neurons, caudal part | medulla oblongata |
| OENr | octavolateralis efferent neurons, rostral part | medulla oblongata |
| P | posterior thalamic nucleus | posterior tuberculum |
| PCN | paracommissural nucleus | diencephalon |
| PGa | anterior preglomerular nucleus | posterior tuberculum |
| PGc | caudal preglomerular nucleus | posterior tuberculum |
| PGl | lateral preglomerular nucleus | posterior tuberculum |
| PGm | medial preglomerular nucleus | posterior tuberculum |
| PGZ | periventricular gray zone of optic tectum | midbrain |
| PL | perilemniscal nucleus | tegmentum |
| PM | magnocellular preoptic nucleus | diencephalon |
| PMg | gigantocellular part of magnocellular preoptic nucleus | diencephalon |
| PO | posterior pretectal nucleus | pretectum |
| PON | posterior octaval nucleus | medulla oblongata |
| PPa | parvocellular preoptic nucleus, anterior part | diencephalon |
| PPd | periventricular pretectal nucleus, dorsal part | pretectum |
| PPp | parvocellular preoptic nucleus, posterior part | diencephalon |
| PPv | periventricular pretectal nucleus, ventral part | pretectum |
| PSm | magnocellular superficial pretectal nucleus | pretectum |
| PSp | parvocellular superficial pretectal nucleus | pretectum |
| PTN | posterior tuberal nucleus | posterior tuberculum |
| PVO | paraventricular organ | posterior tuberculum |
| R | rostrolateral nucleus | thalamus |
| RT | rostral tegmental nucleus | tegmentum |
| SC | suprachiasmatic nucleus | diencephalon |
| SCO | subcommissural organ | diencephalon |
| SD | dorsal sac | epithalamus |
| SG | subglomerular nucleus | posterior tuberculum |
| SGN | secondary gustatory nucleus | medulla oblongata |
| SO | secondary octaval population | medulla oblongata |
| SR | superior raphe | medulla oblongata |
| SRF | superior reticular formation | medulla oblongata |
| SRN | superior reticular nucleus | medulla oblongata |
| T | tangential nucleus | medulla oblongata |
| TeO | optic tectum | midbrain |
| TGN | tertiary gustatory nucleus | posterior tuberculum |
| TL | longitudinal torus | midbrain |
| TLa | lateral torus | posterior tuberculum |
| TPp | periventricular nucleus of posterior tuberculum | posterior tuberculum |
| TSc | central nucleus of semicircular torus | midbrain |
| TSvl | ventrolateral nucleus of semicircular torus | midbrain |
| Val-g | lateral division of valvula cerebelli, granular layer | cerebellum |
| Val-m | lateral division of valvula cerebelli, molecular layer | cerebellum |
| Vam-g | medial division of valvula cerebelli, granular layer | cerebellum |
| Vam-m | medial division of valvula cerebelli, molecular layer | cerebellum |
| VAO | ventral accessory optic nucleus | pretectum |
| Vc | central nucleus of ventral telencephalon area | telencephalon |
| Vd-dd | dorsal zone of ventral telencephalon | telencephalon |
| Vd-vd | ventral zone of ventral telencephalon | telencephalon |
| Vdd | dorsal most zone of ventral telencephalon | telencephalon |
| Vl | lateral nucleus of ventral telencephalon area | telencephalon |
| VL | ventrolateral thalamic nucleus | thalamus |
| VM | ventromedial thalamic nucleus | thalamus |
| Vp | postcommissural nucleus of ventral telencephalon area | telencephalon |
| Vs | supracommissural nucleus of ventral telencephalon area | telencephalon |
| Vv | ventral nucleus of ventral telencephalon area | telencephalon |
| ZL | zona limitans | thalamus |
| III | oculomotor nerve |  |
| IIIm | oculomotor nucleus | tegmentum |
| IVm | trochlear nucleus | tegmentum |
| Vmd | trigeminal motor nucleus, dorsal part | medulla oblongata |
| Vmn | mesencephalic nucleus of the trigeminal nerve | medulla oblongata |
| Vmv | trigeminal motor nucleus, ventral part | medulla oblongata |
| Vsm | primary sensory trigeminal nucleus | medulla oblongata |
| VImc | caudal abducens nerve motor nucleus | medulla oblongata |
| VImr | rostral abducens nerve motor nucleus | medulla oblongata |
| VIILo | facial lobe | medulla oblongata |
| VIIm | facial motor nucleus | medulla oblongata |
| IXLo | glossopharyngeal lobe | medulla oblongata |
| IXm | glossopharyngeal nerve motor nucleus | medulla oblongata |
| XLo | vagal lobe | medulla oblongata |
| Xm | vagal motor nucleus | medulla oblongata |
| UnkD | unknown diencephalon | pretectum |
| UnkMS | unknown mesencephalon | midbrain |
| UnkR | unknown rhombencephalon | medulla oblongata |
| UnkSC | unknown spinal cord | spinal cord |
| UnkVT | unknown ventral telencephalon | telencephalon |

Table S2. Architecture of CNN for detecting cytoplasmic and nuclear *cfos* stained cells.

| Layer (in order) | Output shape |
| --- | --- |
| Input layer | (None, 11, 11, 11, 1) |
| 3D convolution | (None, 11, 11, 11, 64) |
| ReLU | (None, 11, 11, 11, 64) |
| 3D max pooling | (None, 6, 6, 6, 64) |
| Batch normalization | (None, 6, 6, 6, 64) |
| 3D convolution | (None, 6, 6, 6, 64) |
| ReLu | (None, 6, 6, 6, 64) |
| 3D max pooling | (None, 3, 3, 3, 64) |
| Batch normalization | (None, 3, 3, 3, 64) |
| 3D convolution | (None, 3, 3, 3, 128) |
| ReLu | (None, 3, 3, 3, 128) |
| 3D max pooling | (None, 2, 2, 2, 128) |
| Batch normalization | (None, 2, 2, 2, 128) |
| 3D convolution | (None, 2, 2, 2, 256) |
| ReLu | (None, 2, 2, 2, 256) |
| 3D max pooling | (None, 1, 1, 1, 256) |
| Batch normalization | (None, 1, 1, 1, 256) |
| Global average pooling | (None, 256) |
| Dense | (None, 512) |
| ReLu | (None, 512) |
| Dropout | (None, 512) |
| Dense | (None, 1) |
| Activation | (None, 1) |
