## Supplemental File 1: Bench protocol for "Whole-brain mapping in adult zebrafish and identification of a novel tank test functional connectome"

### Bench protocol for whole-brain mapping in zebrafish

#### Reagents (Buffers and Solutions)

**4% PFA**

- 4% PFA
- 1X PBS
- Store at 4C

For 40 mL of solution

- 10 mL of 16% PFA
- 4 mL of 10X PBS
- 26 mL of ultrapure $H_2O$

**5% $H_2O_2$ (Hydrogen peroxide)**

- 5% $H_2O_2$ in MeOH
- Make fresh for each use

For 10 mL of solution

- 3.33 mL of 30% $H_2O_2$
- 6.66 mL of 100% MeOH

**PBS-T**

- 1x PBS
- 0.1% Tween-20
- Store at RT

For 100 mL of solution

- 10 mL of 10X PBS
- 1 mL of 10% Tween-20
- Fill to 100 mL with ultrapure $H_2O$

**5X SSCT Buffer**

- 5X SSC buffer
- 0.1% Tween-20
- Store at RT

For 100 mL of solution

- 5 mL of 20X SSC buffer
- 1 mL of 10% Tween-20
- Fill to 100 mL with ultrapure $H_2O$

**0.25%v/v Acetic anhydride**

- 0.25% Acetic anhydride
- Store at RT

For 50 mL of solution

- 129 µL of 97% Acetic anhydride
- Fill to 50 mL with ultrapure $H_2O$

**Amplification Buffer**

- 5X SSC
- 0.1% Tween-20
- 10% Dextran sulfate
- Store at RT

For 50 mL of solution

- 12.5 mL of 20X SSC buffer
- 500 µL of 10% Tween-20
- 10 mL of 50% dextran sulfate
- Fill to 50 mL with ultrapure $H_2O$

Reagents and supplies

**DNAse/RNAse free 0.5 mL /1.5 mL centrifuge tubes**

**10X PBS (ThermoFisher Cat. #70011-044)**

**Ultrapure H2O (cytiva Cat. # SH30538.03)**

**16% PFA ampoules (Electron Microscopy Sciences Cat.# 15710)**

**10% Tween 20 (Promega Cat.# H5152)**

**20X SSC buffer (Life Technologies Cat # 15557-044)**

**97% Acetic anhydride (ThermoFisher Cat. #A10-500)**

**Dextran sulfate 50% solution (EMD Millipore Corp Cat # S4031))**

**30 % Hydrogen peroxide solution $H_2O_2$ (Sigma-Aldrich Cat. # 216763)**

**100% Methanol (MeOH) (Sigma-Aldrich Cat. # 179337)**

**Dichloromethane (DCM) (Sigma-Aldrich Cat. # 270997)**

**Dibenzyl Ether (DBE) (Sigma-Aldrich Cat. # 108014)**

**Probe mixture *for target of interest, e.g. cfos/fosab* (Molecular Instruments)**

**Probe hybridization buffer (Molecular Instruments)**

**Probe Wash buffer (Molecular Instruments)**

**Hairpins (Molecular Instruments)**
***Note: The hairpin initiator sequence MUST correspond to the initiator sequence present on the detection probes (e.g. B1). There should be two sets of hairpins (e.g. H1B1 and H2B1)***

**Note: We purchase probe hybridization and wash buffer from Molecular Instruments. The recipes for these solutions can be found in the supplemental material of Choi et al (2014) https://doi.org/10.1021/nn405717p**

#### Sample Preparation

1. Record behavior, then anesthetize/euthanize animals in ice water slurry for at least 1 min.
2. Decapitate fish using a sharp blade and wash head in 1X PBS for 30-60 seconds to allow blood to drain.
3. Fix head in 4% PFA at 4°C overnight.
    - We use 1.5 mL tubes filled with 1 mL PFA.
4. Carefully dissect brains out in cold 1X PBS and place in 0.5 mL tubes containing 1X PBS and proceed to pretreatment step of tissue clearing.
    - Tissue can be held in 1X PBS in 0.5 mL tubes on ice while all brains are dissected.

***Note: Perform all steps on ice***

#### Pretreatment of Samples

1. Wash sample 3 x 30 min in 1X PBS at RT with shaking.
2. Dehydrate samples using MeOH gradient (20%, 40%, 60%, 80%, 100%) for 30 min each, without shaking.
3. Perform extra 100% MeOH wash at 4°C for 1hr.

***Note: Make and chill bleach solution at this step so it's cold for the follow up step.***

4. Bleach samples in 5% $H_2O_2$ in MeOH (1 volume 30% $H_2O_2$ to 5 volumes MeOH) overnight at 4°C.
5. Rehydrate samples using MeOH gradient (80%, 60%, 40%, 20%) for 30 min each at RT without shaking.
6. Wash samples with 1X PBS for 1hr, then twice in PBS-T (1hr, then 3hr).
7. Equilibrate samples in 5X SSCT buffer overnight. The samples are now ready for the preparation phase of *in situ* HCR.

#### Preparation Phase for *in-situ* HCR

1. Acetylate samples with 0.25% v/v acetic anhydride for 30 mins at RT.
2. Wash samples 3x with ultrapure water for 5 min each at RT.

#### Detection Phase for *in-situ* HCR

1. Wash samples in probe hybridization buffer for 15 min at RT.
2. Remove buffer, then incubate in probe hybridization buffer for 1 hr at 37°C with shaking.

***Note: Hybridization buffer must be warmed to room temperature before use (~1 hr).***
***Note: During this step, warm 600 µL per sample of probe hybridization buffer to 37°C.***

**Caution:**

- ***Probe hybridization solution contains formamide. This is a hazardous chemical so these steps should be done in the fume hood. All waste should be disposed of as hazardous waste.***
- ***Brain tissue floats in probe hybridization buffer, make sure not to poke them while changing the solution.***

***Note: Before pipetting the probes, thaw them on ice (~30 minutes), vortex after thawing, and spin down briefly.***

3. Prepare probe hybridization solution: 2 µL of 1 µM probe stock for each 600 µL pre-warmed buffer at 37°C.
4. Add 600 µL probe hybridization solution to each sample and incubate for 48-60 hr at 37°C with shaking.

***Note: After incubation is finished, remove as much probe wash buffer as you need and pre-warm to 37°C before adding to samples.***

5. Wash samples 3x 60 min with pre-warmed probe wash buffer at 37°C with shaking.

***Caution: Probe wash solution contains formamide so these steps should be performed in a fume hood. Waste should be disposed of as hazardous waste.***

6. Wash samples 2 x 60 min with 5X SSCT at RT with shaking.
    - Samples are now ready for the amplification stage.

#### Amplification Phase for In-situ HCR

1. Incubate samples for 1 hr in 600 µL amplification buffer at RT with shaking.

***Note: Equilibrate amplification buffer to room temperature prior to use.***
***Note: After starting step 1, begin to prepare the hairpins.***

2. Before beginning, warm up a thermal cycler to 95°C.
3. Calculate the amount of each hairpin needed for your final concentration
    - We use 7.5 pmol in 125 µL of H1 and H2 for each sample (i.e., 2.5 µL of 3 µM hairpin solution). This is the same concentration recommended by molecular instruments but saves money by using a smaller volume.
4. Thaw hairpins in the dark on ice/4°C.
5. Vortex hairpins such that the blue dye concentrated at the bottom of the tube is completely mixed in the solution. Then briefly spin down tubes containing hairpins before opening (on small centrifuge).
6. Pipette hairpins (e.g. 2.5 µL of H1 and H2 each) per sample into PCR tubes for each hairpin ***(this is important, separate tubes, one tube for H1 hairpin and other tube for H2 hairpins!)***
7. Heat hairpins to 95°C in PCR machine for 90 sec and cool to RT in a dark drawer for 30 min.
8. Add both hairpins to amplification buffer at room temperature and mix with pipette.
9. Add hairpin/amplification buffer solution to each sample
    - We add 130 µL per sample (125 µL buffer + 2.5 µl of H1 +2.5 µl of H2).
10. Incubate for two days (48 hours) in the dark at room temperature.
11. Wash samples with 5X SSCT 5 x 60 min in the dark at RT.
    - Leave last wash overnight before clearing.

#### Clearing

1. Dehydrate samples using a MeOH gradient of 20%, 40%, 60%, 80%, and 100% for 1hr at RT.
2. Wash an additional time in 100% MeOH for 1hr at RT.
3. Samples can be left in MeOH overnight or can be continued to the next step.
4. Incubate samples in DCM/MeOH (66%/33%) mix at RT for 3 hours.

***Caution:***

- ***This procedure must be performed in the fume hood. MeOH and DCM waste must be disposed of as hazardous waste.***
- ***When working with DCM you must wear two pairs of 8 mil nitrile gloves. DCM will quickly penetrate typical 4 mil gloves.***
- ***To obtain DCM, use 10 mL syringe and 26-gauge needle. Carefully extract the solution. It can be difficult to get the volume exactly right; it is okay to get an approximate volume of DCM.***

5. Carefully remove DCM/MeOH solution from sample tubes and was samples 2 x 15 min each in 100% DCM at RT.
    - Samples often float in DCM so be extra careful not to poke samples with your pipette tip. It is okay to leave a little bit of DCM/MeOH in tube. Better to leave extra than damage the brain. However, for the last DCM wash, as much should be removed as possible before adding in DBE.

6. Carefully remove as much DCM as possible and incubate samples in DBE at RT in the dark. Samples should remain in DBE for at least 24 hr to ensure adequate clearing before imaging.

***Note: To add DBE to samples, pour DBE from larger bottle into a clean small glass beaker (less than 50 mL). Pipette approximately 600 µL of DBE from the glass beaker into each tube. Once done, dispose of the extra DBE by dumping it into a hazardous waste disposal container. Wipe the beaker with Kimwipes and then wash it with ethanol. Leave the beaker in the fume hood overnight for the ethanol to evaporate before sending the beaker to be washed.***

***Caution: This procedure must be performed in the fume hood. DBE and DCM waste must be disposed of as hazardous waste. Under no circumstances is DBE to be washed down the drain as it is highly toxic to aquatic organisms***.

#### Imaging:

To perform imaging, mount the cleared samples on the sample holder using an ultraviolet-curing resin (Norland Optical Adhesive 61, refractive index 1.56) that matches the refractive index of the imaging solution, DBE. We use a Miltenyi LaVision BioTec UltraMicroscope II light sheet system with Imspector software for image acquisition, and terastitcher for stitching.

#### Cell quantification using CellFinder:

To automatically detect and quantify *cfos* positive cells, we use the Python-based software, CellFinder. It comprises two steps: (1) cell detection and (2) cell classification.

***Note: To setup CellFinder and its requirements on your system. Follow the steps mentioned in the Brainglobe documentation.***
***https://brainglobe.info/documentation/setting-up/index.html***

**Steps:**

1. The initial step of cell detection requires parameter optimization, like soma diameter, filter size for cell intensity and threshold value, to identify all potential cell candidates. Using the Linux terminal, run the following cell detection command to detect all the possible cell candidates. You will need to input your x,y, and z resolutions from your images.

```
cellfinder -s path/to/folder/signal/channel/cfos/ -b
/path/to/folder/background/channel/AF -o path/to/output1 -v
z_res x_res y_res --orientation sal --no-register --no-
classification --soma-diameter 5 --threshold 3 --ball-xy-size 2
```

```
--ball-z-size 7 --ball-overlap-fraction 0.77 --log-sigma-size
0.1 --save-csv --batch-size 64 --epochs 100
```

***Note: Several of these parameters will need to be adjusted for your specific settings, such as the resolution (after -v) and the orientation. See the CellFinder documents for more details.***

***Pro tip: For running multiple files in the folder rather than doing one file at a time. Run the command using for loop.***

```
for f in /path_to_the_folder/folder/*; do cellfinder -s
/$f/cfos/ -b /$f/AF/ -o /$f/wholebrain -v z_res x_res y_res --
orientation sal --no-register --no-classification --soma-
diameter 5 --threshold 3 --ball-xy-size 2 --ball-z-size 7 --
ball-overlap-fraction 0.77 --log-sigma-size 0.1 --save-csv; done
```

2. After detecting all the possible cell candidates, use a python script to remove the overlapped annotated cell candidates ***(https://github.com/KenneyLab/RajputEtAl_2024_Whole_brain_mapping/blob/main/remove_overlaps.py)***. Optimization of filter size and cell number per split will be necessary. We used the following command to run the script.

```
python /path/to/file/remove_overalps.py --
file=output1/points/cells.xml --filter=9 --cellsplits=12000 --
x_res=x.xxx --y_res=y.yyy --z_res=z.zzz
```

3. After detecting cell candidates, use napari to visualize the filtered cells.xml file. Thereafter, either train a network according to the requirement or utilize the default pre-trained network provided with the CellFinder package.  We suggest training a new network and to incorporate images from a variety of brains to ensure you capture all the variability in imaging of your samples. This may take a few rounds of training and testing to get right.
4. To train the network from scratch manually label cells and non-cells. This is done by loading the filtered cells.xml on to napari. Then from tabs click on “Plugins”, then select the curation option. Save the manually annotated data as .yml files and then train the network using these .yml files. More details can be found in the CellFinder documentation Use the following cellfinder train command:

```
Cellfinder_train -y path/to/labels_brain1.yml
path/to/labels_brain2.yml path/to/labels_brain3.yml -o
/trained_network --batch-size 64 --epochs 100 --no-save-
checkpoints --save-progress
```

5. Finally, apply the trained network to all the experimental brains to classify the detected cell candidates into cells and non-cells. Use the following cell classification command with parameters determined from your testing above:

```
cellfinder -s /path/to/folder/signal/channel/cfos/ -b
/path/to/folder/background/channel/AF/ -o path/to/output1 -v
z_res x_res y_res --orientation sal --no-register --no-detection
--soma-diameter 5 --threshold 3 --ball-xy-size 2 --ball-z-size 7
--ball-overlap-fraction 0.77 --log-sigma-size 0.1 --save-csv --
trained-model /trained_network/model.h5
```

#### Registration

For automated segmentation of brain regions in new images using a brain atlas we register to an existing atlas using advanced normalization tools (ANTs). Registration is essential to align one image with another, ensuring that corresponding voxels represent the same structures in both images. For optimal alignment between brain images and the atlas, you will need to optimize four key parameters of the diffeomorphic algorithm: cross-correlation, gradient step, update field variance in voxel space, and total field variance in voxel space.

***Note: Range of parameters we explored to optimize the diffeomorphic algorithm.***
***Cross-correlation (2 – 4, step of 1)***
***Gradient step (0.1 – 0.4, step of 0.1)***
***Update field variance (2 - 8, step of 2)***
***Total field variance (0, 0.5)***

**Steps:**

1. Images need to be down sampled to match the resolution in the atlas you are using. AZBA is at 4 µm isotropic. Brainreg will automatically down sample your image to match the atlas.

```
brainreg /background_channel /output/folder/ -v
your_data_voxel_size --orientation your_data_orientation --atlas
azba_zfish_4um --debug
```

For example, we used:

```
brainreg /AF /REG/ -v 3.990 0.943 0.943 --orientation sal --
atlas azba_zfish_4um --debug
```

***Note: Downsampled images can also be generated using c3d through a multi-step process. First step is tiling, the TIFF into a nifti stack. Then assigning the***

***appropriate dimensions using the '-spacing' command, and then resampling using '-resample'***

```
c3d *.tif -tile z -o volume.nii.gz
c3d volume.nii -spacing z_vox x x_vox x y_vox -o output.nii
c3d output.nii -resample-mm z_vox x x_vox x y_vox -o output2.nii
```

2. Then reorient the downsampled images to "ASL" using the convert 3D i.e. c3d image processing program. Use the following command line to match AZBA.

```
c3d input_downsampled.nii -orient ASL -o output_downsampled.nii
```

***Note: Downsampled images can be reoriented using ITK snap. To do so, open ITK snap load your downsampled.nii file. Then from the tabs select the "Tools" option. Select the reorient option and change new orientation to "ASL". Save the downsampled.nii file.***

3. For creating the average template select a handful of your best brain images (~10). Use brains that are dissected without major nicks or pokes.

4. For registration, out of the selected brain images, use one image as the template (fixed image) and register rest images on to the template (moving images). This is done by using the following ANTS command (replace parameters with those you optimized for your samples).

```
antsRegistration --dimensionality 3 --float 1 -o
[${AF_sample_1_for_avg_},${ AF_sample_1_for_avg-warped}] --
interpolation WelchWindowedSinc -u 0 -r
[${template.nii},${AF_sample_1.nii},1] -t Rigid[0.1] -m
MI[${template.nii},${AF_sample_1.nii},1,32,Regular,0.25] -c [200
x 200 x 200 x 0,1e-8,10] --shrink-factors 12x8x4x2 --smoothing-
sigmas 4x3x2x1vox -t Affine[0.1] -m
MI[${template.nii},${AF_sample_1.nii}, 1,32,Regular,0.25] -c
[200 x 200 x 200 x 0,1e-8,10] --shrink-factors 12x8x4x2 --
smoothing-sigmas 4x3x2x1vox -t SyN[0.3,4,0] -m
CC[${template.nii},${AF_sample_1.nii}, 1,3] -c [200 x 200 x 200
x 200, 1e-6,10] --shrink-factors 12x8x4x2 --smoothing-sigmas
4x3x2x1vox --verbose 1
```

5. Use the output generated from the first registration step to create the averages of the samples. To generate the average template, use the following ANTs AverageImages command:

```
AverageImages 3 avg_template.nii.gz 1 AF_sample_1_for_avg-
warped.nii.gz, AF_sample_2_for_avg-warped.nii.gz…
```

6. Following this, register AZBA (moving image) to the average template (fixed image). That is bringing AZBA into the same space of average template using the ANTs command:

```
antsRegistration --dimensionality 3 --float 1 -o
[${AZBA_to_avg_temp_},${AZBA_to_avg_temp-warped}] --
interpolation WelchWindowedSinc -u 0 -r
[${avg_template.nii.gz},${AZBA/20180628_AF_average.nii.gz},1] -t
Rigid[0.1] -m
MI[${avg_template.nii.gz},${AZBA/20180628_AF_average.nii.gz},1,3
2,Regular,0.25] -c [200 x 200 x 200 x 0,1e-8,10] --shrink-
factors 12x8x4x2 --smoothing-sigmas 4x3x2x1vox -t Affine[0.1] -m
MI[${avg_template.nii.gz},${AZBA/20180628_AF_average.nii.gz},
1,32,Regular,0.25] -c [200 x 200 x 200 x 0,1e-8,10] --shrink-
factors 12x8x4x2 --smoothing-sigmas 4x3x2x1vox -t SyN[0.3,4,0] -
m CC[${avg_template.nii.gz},${AZBA/20180628_AF_average.nii.gz},
1,3] -c [200 x 200 x 200 x 200, 1e-6,10] --shrink-factors
12x8x4x2 --smoothing-sigmas 4x3x2x1vox --verbose 1
```

7. Transform AZBA segmentation into the space of average template (reference image) using the following ANTs command:

```
antsApplyTransforms -d 3 --float -n NearestNeighbor -i
/AZBA/2021-08-22_AZBA_segmentation.nii.gz -r avg_template.nii.gz
-o AZBA_to_avg_temp_transformed.nii.gz -t
AZBA_to_avg_temp_1Warp.nii.gz -t
AZBA_to_avg_temp_0GenericAffine.mat
```

8. Finally, register the average template image (moving image) onto all individual downsampled autofluorescence images of the experimental brains (fixed image) using the following ANTs command:

```
antsRegistration --dimensionality 3 --float 1 -o
[${AF_sample_},${AF_sample-warped}] --interpolation
WelchWindowedSinc -u 0 -r
[${AF_sample.nii},${avg_template.nii.gz },1] -t Rigid[0.1] -m
MI[${AF_sample.nii},${ avg_template.nii.gz },1,32,Regular,0.25]
-c [200 x 200 x 200 x 0,1e-8,10] --shrink-factors 12x8x4x2 --
smoothing-sigmas 4x3x2x1vox -t Affine[0.1] -m
MI[${AF_sample.nii},${ avg_template.nii.gz },1,32,Regular,0.25]
-c [200 x 200 x 200 x 0,1e-8,10] --shrink-factors 12x8x4x2 --
smoothing-sigmas 4x3x2x1vox -t SyN[0.3,4,0] -m
CC[${AF_sample.nii},${ avg_template.nii.gz },1,3] -c [200 x 200
x 200 x 200, 1e-6,10] --shrink-factors 12x8x4x2 --smoothing-
sigmas 4x3x2x1vox --verbose 1
```

9. Transform average template segmentation into the space of all experimental brains by using the ANTs transformation command:

```
antsApplyTransforms -d 3 --float -n NearestNeighbor -i
AZBA_to_avg_temp_transformed.nii.gz -r AF_sample.nii -o
AF_sample_transformed.nii.gz -t AF_sample_1Warp.nii.gz -t
AF_sample_0GenericAffine.mat
```

***Pro tip: To run the above steps for multiple brain ID's use this command line in terminal to run loop through all ID's.***

***To run the loop from terminal, make ".sh" file (e.g. registration.sh) with the ANTS registration command written in it. Additionally, make one .txt file (e.g. brain_id_list.txt) with the list of all brain ID's.***

***Open a terminal, then navigate to the folder where both the files (.sh and .txt) and AF_downsampled.nii brain images for each corresponding brain IDs are present. Run this command on terminal:***

```
cat brain_ID_list.txt | while read l; do sh ANTS_regsitartion.sh
“$l” *.nii $l; done
```

Your final output will be a .nii file containing the segmentation for each individual brain. This is then combined with the output from cellfinder for each individual brain. One way to combine these files is to use R and the RNifti package. The code for this can be found on our github page:
https://github.com/KenneyLab/RajputEtAl_2024_Whole_brain_mapping/
